# Testing pleiotropy vs. separate QTL in multiparental populations

**DOI:** 10.1101/550939

**Authors:** Frederick J. Boehm, Elissa J. Chesler, Brian S. Yandell, Karl W. Broman

**Affiliations:** Department of Statistics, University of Wisconsin-Madison, Madison, Wisconsin 53706; The Jackson Laboratory, Bar Harbor, Maine 04609; Department of Horticulture, University of Wisconsin-Madison, Madison, Wisconsin 53706; Department of Biostatistics and Medical Informatics, University of Wisconsin-Madison, Madison, Wisconsin 53706

**Keywords:** Quantitative trait locus, pleiotropy, multivariate analysis, linear mixed effects models, systems genetics

## Abstract

The high mapping resolution of multiparental populations, combined with technology to measure tens of thousands of phenotypes, presents a need for quantitative methods to enhance understanding of the genetic architecture of complex traits. When multiple traits map to a common genomic region, knowledge of the number of distinct loci provides important insight into the underlying mechanism and can assist planning for subsequent experiments. We extend the method of Jiang and Zeng (1995), for testing pleiotropy with a pair of traits, to the case of more than two alleles. We also incorporate polygenic random effects to account for population structure. We use a parametric bootstrap to determine statistical significance. We apply our methods to a behavioral genetics data set from Diversity Outbred mice. Our methods have been incorporated into the R package `qtl2pleio`.

---

Complex trait studies in multiparental populations present new challenges in statistical methods and data analysis. Among these is the development of strategies for multivariate trait analysis. The joint analysis of two or more traits allows one to address additional questions, such as whether two traits share a single pleiotropic locus.

Previous research addressed the question of pleiotropy vs. separate QTL in two-parent crosses. Jiang and Zeng (1995) developed a likelihood ratio test for pleiotropy vs. separate QTL for a pair of traits. Their approach assumed that each trait was affected by a single QTL. Under the null hypothesis, the two traits were affected by a common QTL, and under the alternative hypothesis the two traits were affected by distinct QTL. Knott and Haley (2000) used linear regression to develop a fast approximation to the test of Jiang and Zeng (1995), while Tian *et al*. (2016) used the methods from Knott and Haley (2000) to dissect QTL hotspots in a F_2_ population.

Multiparental populations, such as the Diversity Outbred (DO) mouse population (Churchill *et al*. 2012), enable high-precision mapping of complex traits (de Koning and McIntyre 2014). The DO mouse population began with progenitors of the Collaborative Cross (CC) mice (Churchill *et al*. 2004) Each DO mouse is a highly heterozygous genetic mosaic of alleles from the eight CC founder lines. Random matings among non-siblings have maintained the DO population for more than 23 generations (Chesler *et al*. 2016).

Several limitations of previous pleiotropy vs. separate QTL tests prevent their direct application in multiparental populations. First, multiparental populations can have complex patterns of relatedness among subjects, and failure to account for these patterns of relatedness may lead to spurious results (Yang *et al*. 2014). Second, previous tests allowed for only two founder lines (Jiang and Zeng 1995). Finally, Jiang and Zeng (1995) assumed that the null distribution of the test statistic follows a chi-square distribution.

We developed a pleiotropy vs. separate QTL test for two traits in multiparental populations. Our test builds on research that Jiang and Zeng (1995), Knott and Haley (2000), Tian *et al*. (2016), and Zhou and Stephens (2014) initiated. Our innovations include the accommodation of *k* founder alleles per locus (compared to the traditional two founder alleles per locus) and the incorporation of multivariate polygenic random effects to account for relatedness. Furthermore, we implemented a parametric bootstrap test to assess statistical significance (Efron 1979; Tian *et al*. 2016). We focus on the case that two traits are measured in the same set of subjects (Design I in the notation of Jiang and Zeng (1995)).

Below, we describe our likelihood ratio test for pleiotropy vs. separate QTL. In simulation studies, we find that it is slightly conservative, and that it has power to detect two separate loci when the univariate LOD peaks are strong. We further illustrate our approach with an application to data on a pair of behavior traits in a population of 261 DO mice (Logan *et al*. 2013; Recla *et al*. 2014).

## Methods

Our strategy involves first identifying two traits that map to a common genomic region. We then perform a two-dimensional, two-QTL scan over the genomic region, with each trait affected by one QTL of varying position. We identify the QTL position that maximizes the likelihood under pleiotropy (that is, along the diagonal where the two QTL are at a common location), and the ordered pair of positions that maximizes the likelihood under the model where the two QTL are allowed to be distinct. The logarithm of the ratio of the two likelihoods is our test statistic. We determine statistical significance with a parametric bootstrap.

### Data structures

The data consist of three objects. The first is an *n* by *k* by *m* array of allele probabilities for *n* subjects with *k* alleles and *m* marker positions on a single chromosome [derived from the observed SNP genotype data by a hidden Markov model; see Broman *et al*. (2019)]. The second object is an *n* by 2 matrix of phenotype values. Each column is a phenotype and each row is a subject. The third object is an *n* by *c* matrix of covariates, where each row is a subject and each column is a covariate.

One additional object is the genotype-derived kinship matrix, which is used in the linear mixed model to account for population structure. We are focusing on a defined genomic interval, and we prefer to use a kinship matrix derived by the “leave one chromosome out” (LOCO) method (Yang *et al*. 2014), in which the kinship matrix is derived from the genotypes for all chromosomes except the chromosome under test.

### Statistical Models

Focusing on a pair of traits and a particular genomic region of interest, the next step is a two-dimensional, two-QTL scan (Jiang and Zeng 1995). We consider two QTL with each affecting a different trait, and consider all possible pairs of locations for the two QTL. For each pair of positions, we fit the multivariate linear mixed effects model defined in Equation 1. Note that we have assumed an additive genetic model throughout our analyses, but extensions to design matrices that include dominance are straightforward.

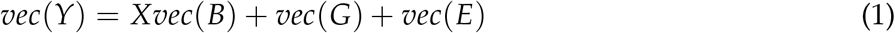

where *Y* is the *n* by 2 matrix of phenotypes values; *X* is a 2*n* by 2(*k* + *c*) matrix that contains the *k* allele probabilities for the two QTL positions and the *c* covariates in diagonal blocks; *B* is a (*k* + *c*) by 2 matrix of allele effects and covariate effects; *G* is a *n* by 2 matrix of random effects; and *E* is a *n* by 2 matrix of random errors. *n* is the number of mice. The ‘vec’ operator stacks columns from a matrix into a single vector. For example, a 2 by 2 matrix inputted to ‘vec’ results in a vector with length 4. Its first two entries are the matrix’s first column, while the third and fourth entries are the matrix’s second column.

We also impose distributional assumptions on *G* and *E*:

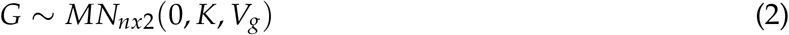

and

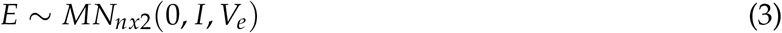

where *MN*_*nx*2_ (0, *V_r_*, *V_c_*) denotes the matrix-variate (*n* by 2) normal distribution with mean being the *n* by 2 matrix with all zero entries and row covariance *V_r_* and column covariance *V_c_*. We assume that *G* and *E* are independent.

### Parameter inference and log likelihood calculation

Inference for parameters in multivariate linear mixed effects models is notoriously difficult and can be computationally intense (Meyer 1989, 1991). Thus, we estimate *V_g_* and *V_e_* under the null hypothesis of no QTL, and then take them as fixed and known in our two-dimensional, two-QTL genome scan. We use restricted maximum likelihood methods to fit the model:

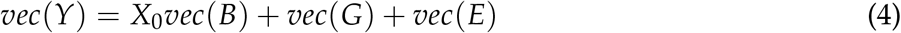

where *X*_0_ is a 2*n* by 2(*c* + 1) matrix whose first column of each diagonal block in *X*_0_ has all entries equal to one (for an intercept); the remaining columns are the covariates.

We draw on our R implementation (Boehm 2018) of the GEMMA algorithm for fitting a multivariate linear mixed effects model with expectation-maximization (Zhou and Stephens 2014). We use restricted maximum likelihood fits for the variance components *V_g_* and *V_e_* in subsequent calculations of the generalized least squares solution 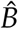.

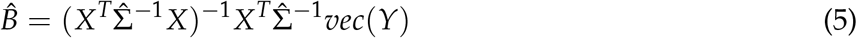

where

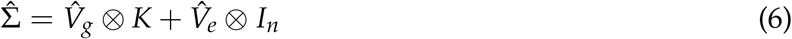

where ⊗ denotes the Kronecker product, *K* is the kinship matrix, and *I_n_* is a n by n identity matrix. We then calculate the log likelihood for a normal distribution with mean 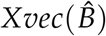 and covariance 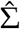 that depends on our estimates of *V_g_* and *V_e_* (Equation 6).

### Pleiotropy vs. separate QTL hypothesis testing framework

Our test applies to two traits considered simultaneously. Below, *λ*_1_ and *λ*_2_ denote putative locus positions for traits one and two. We quantitatively state the competing hypotheses for our test as:

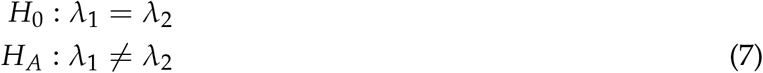

Our likelihood ratio test statistic is:

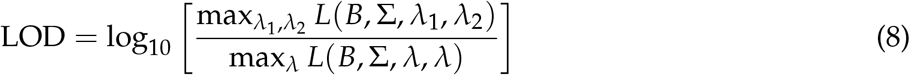

where *L* is the likelihood for fixed QTL positions, maximized over all other parameters. The denominator concerns the likelihood for the null hypothesis of pleiotropy, where *λ* = *λ*_1_ = *λ*_2_.

### Visualizing profile LOD traces

The output of the above analysis is a two-dimensional log10 likelihood surface. To visualize these results, we followed an innovation of Zeng *et al*. (2000) and Tian *et al*. (2016), and plot three traces: the results along the diagonal (corresponding to the null hypothesis of pleiotropy), and then the profiles derived by fixing one QTL’s position and maximizing over the other QTL’s position.

We define the LOD score for our test:

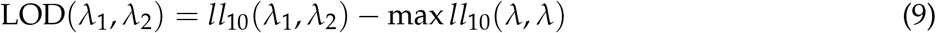

where *ll*_10_ denotes log_10_ likelihood.

We follow Zeng *et al*. (2000) and Tian *et al*. (2016) in defining profile LOD by the equation

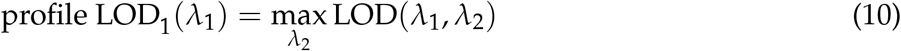

We define profile LOD_2_(*λ*_2_) analogously. The profile LOD1 and profile LOD_2_ traces have the same maximum value, which is non-negative and gives the overall LOD test statistic.

We construct the pleiotropy trace by calculating the log-likelihoods for the pleiotropic models at every position.

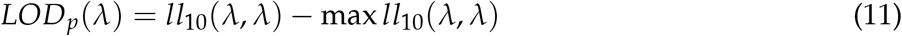

By definition, the maximum value for this pleiotropy trace is zero.

### Bootstrap for test statistic calibration

We use a parametric bootstrap to determine statistical significance (Efron 1979). While Jiang and Zeng (1995) used quantiles of a chi-squared distribution to determine p-values, this does not account for the two-dimensional search over QTL positions. We follow the approach of Tian *et al*. (2016), and identify the maximum likelihood estimate of the QTL position under the null hypothesis of pleiotropy. We then use the inferred model parameters under that model and with the QTL at that position to simulate bootstrap data sets according to the model in equations 1–3. For each of *b* bootstrap data sets, we perform a two-dimensional QTL scan (over the genomic region of interest) and derive the test statistic value. We treat these *b* test statistics as the empirical null distribution, and calculate a p-value as the proportion of the *b* bootstrap test statistics that equal or exceed the observed one, with the original data, 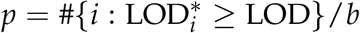 where 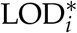 denotes the LOD score for the ith bootstrap replicate and LOD is the observed test statistic.

### Data & Software Availability

Our methods have been implemented in an R package, `qtl2pleio`, available at GitHub: https://github.com/fboehm/qtl2pleio

Custom R code for our analyses and simulations are at GitHub: https://github.com/fboehm/qt2pleio-manuscript-clean

The data from Recla *et al*. (2014) and Logan *et al*. (2013) are available at the Mouse Phenome Database: https://phenome.jax.org/projects/Chesler4 and https://phenome.jax.org/projects/Recla1.

They are also available in R/qtl2 format at https://github.com/rqtl/qtl2data.

## Simulation studies

We performed two types of simulation studies, one for type I error rate assessment and one to characterize the power to detect separate QTL. To simulate traits, we specified *X, B, V_g_, K*, and *V_e_* matrices (Equations 1–3). For both we used the allele probabilities from a single genomic region derived empirically from data for a set of 479 Diversity Outbred mice from Keller *et al*. (2018).

### Type I error rate analysis

To quantify type I error rate (*i.e*., false positive rate), we simulated 400 pairs of traits for each of eight sets of parameter inputs (Table 1). We used a 2^3^ factorial experimental design with three factors: allele effects difference, allele effects partitioning, and genetic correlation, *i.e*., the off-diagonal entry in the 2 by 2 matrix *V_g_*.

**Table 1.** Type I error rates for all runs in our 2^3^ experimental design. We set (marginal) genetic variances (*i.e*., diagonal elements of *V_g_*) to 1 in all runs. *V_e_* was set to the 2 by 2 identity matrix in all runs. We used allele probabilities at a single genetic marker to simulate traits for all eight sets of parameter inputs. In the column “Allele effects partitioning”, “ABCD:EFGH” means that lines A–D carry one QTL allele while lines E–H carry the other allele. “F:ABCDEGH” means the QTL has a private allele in strain F.

| Run | $\Delta(\text{Allele effects})$ | Allele effects partitioning | Genetic correlation | Type I error rate |
| --- | --- | --- | --- | --- |
| 1 | 6 | ABCD:EFGH | 0 | 0.032 |
| 2 | 6 | ABCD:EFGH | 0.6 | 0.035 |
| 3 | 6 | F:ABCDEGH | 0 | 0.040 |
| 4 | 6 | F:ABCDEGH | 0.6 | 0.045 |
| 5 | 12 | ABCD:EFGH | 0 | 0.038 |
| 6 | 12 | ABCD:EFGH | 0.6 | 0.042 |
| 7 | 12 | F:ABCDEGH | 0 | 0.025 |
| 8 | 12 | F:ABCDEGH | 0.6 | 0.025 |

We chose two strong allele effects difference values, 6 and 12. These ensured that the univariate phenotypes mapped with high LOD scores to the region of interest. For the allele partitioning factor, we used either equally frequent QTL alleles, or a private allele in the CAST strain (F). For the residual genetic correlation (the off-diagonal entry in *V_g_*), we considered the values 0 and 0.6. The marginal genetic variances (*i.e*., the diagonal entries in *V_g_*) for each trait were always set to one.

We performed 400 simulation replicates per set of parameter inputs, and each used *b* = 400 bootstrap samples. For each bootstrap sample, we calculated the test statistic (Equation 8). We then compared the test statistic from the simulated trait against the empirical distribution of its 400 bootstrap test statistics. When the simulated trait’s test statistic exceeded the 0.95 quantile of the empirical distribution of bootstrap test statistics, we rejected the null hypothesis. We observed that the test is slightly conservative over our range of parameter selections (Table 1), with estimated type I error rates < 0.05.

### Power analysis

We also investigated the power to detect the presence of two distinct QTL. We used a 2 × 2 × 5 experimental design, where our three factors were allele effects difference, allele effects partitioning, and inter-locus distance. The two levels of allele effects difference were 1 and 2. The two levels of allele effects partitioning were as in the type I error rate studies, ABCD:EFGH and F:ABCDEGH (Table S1). The five levels of interlocus distance were 0, 0.5, 1, 2, and 3 cM. *V_g_* and *V_e_* were both set to the 2 by 2 identity matrix in all power study simulations.

We simulated 400 pairs of traits per set of parameter inputs. For each simulation replicate, we calculated the likelihood ratio test statistic. We then applied our parametric bootstrap to determine the statistical significance of the results. For each simulation replicate, we used *b* = 400 bootstrap samples. Because the bootstrap test statistics within a single set of parameter inputs followed approximately the same distribution, we pooled the 400 * 400 = 160,000 bootstrap samples per set of parameter inputs and compared each test statistic to the empirical distribution derived from the 160,000 bootstrap samples. However, for parameter inputs with interlocus distance equal to zero, we did not pool the 160,000 bootstrap samples; instead, we proceeded by calculating power (*i.e*., type I error rate, in this case), as we did in the type I error rate study above.

We present our power study results in Figure 1. Power increases as interlocus distance increases. The top two curves correspond to the case where the QTL effects are largest. For each value for the QTL effect, power is greater when the QTL alleles are equally frequent, and smaller when a QTL allele is private to one strain. One can have high power to detect that the two traits have distinct QTL when they are separated by > 1 cM and when the QTL have large effect. We provide example profile LOD plots from the power analysis in Figure S3.

**Figure 1.**
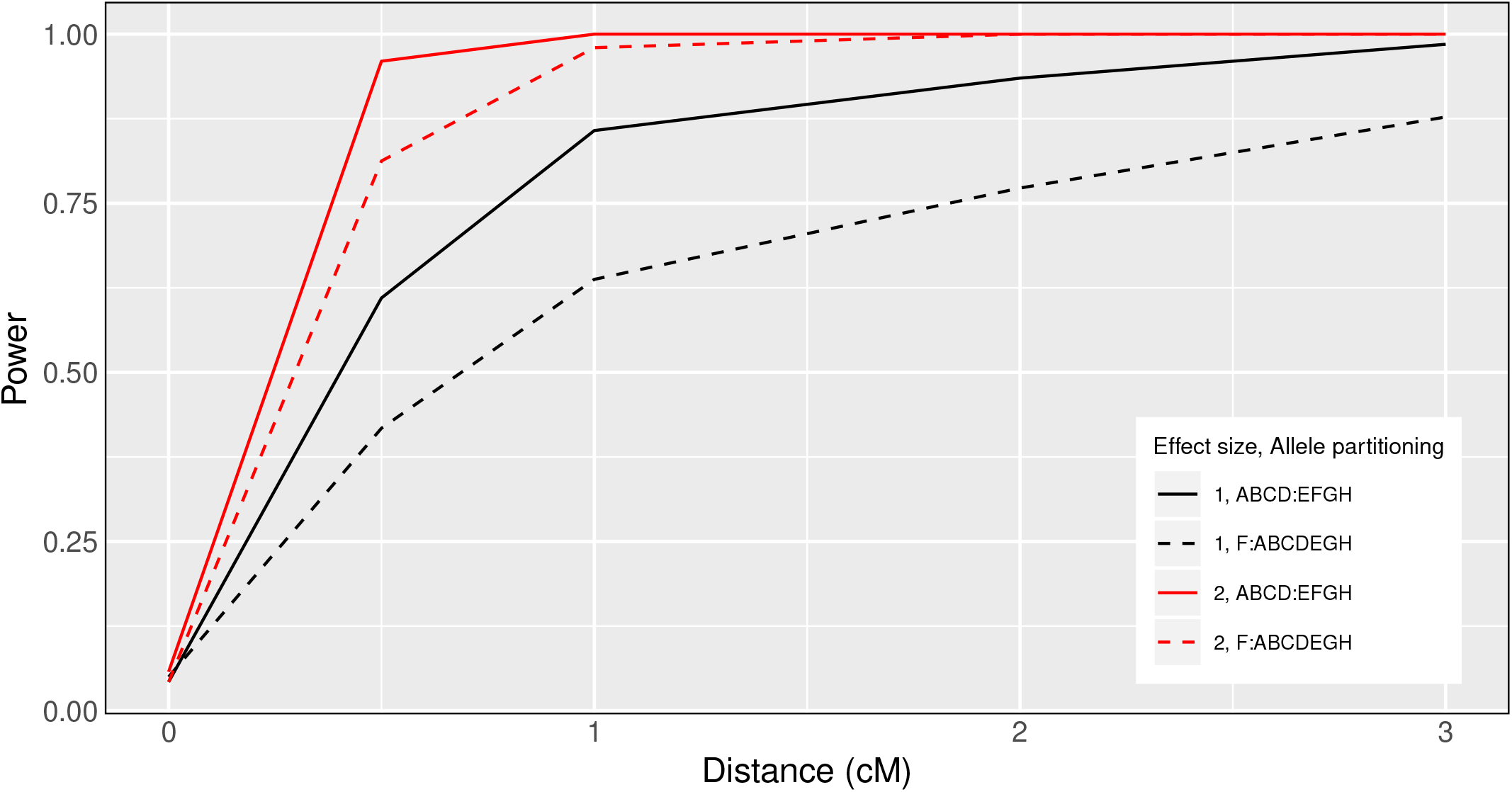
Pleiotropy vs. separate QTL power curves for each of four sets of parameter settings. Factors that differ among the four curves are allele effects difference and allele partitioning. Red denotes high allele effects difference, while black is the low allele effects difference. Solid line denotes the even allele partitioning (ABCD:EFGH), while dashed line denotes the uneven allele partitioning (F:ABCDEGH).

## Application

To illustrate our methods, we applied our test to data from Logan *et al*. (2013) and Recla *et al*. (2014), on 261 DO mice measured for a set of behavioral phenotypes. Recla *et al*. (2014) identified *Hydin* as the gene that underlies a QTL on Chromosome 8 at 57 cM for the “hot plate latency” phenotype (a measure of pain tolerance). The phenotype “percent time in light” in a light-dark box (a measure of anxiety) was measured on the same set of mice (Logan *et al*. 2013) and also shows a QTL near this location, which led us to ask whether the same locus affects both traits. The two traits show a correlation of −0.15 (Figure S1).

QTL analysis with the LOCO method, and using sex as an additive covariate, showed multiple suggestive QTL for each phenotype (Figure S2; Table S2). For our investigation of pleiotropy, we focused on the interval 53–64 cM on Chromosome 8. Univariate analyses showed a QTL in this region for both traits (Figure 2).

**Figure 2.**
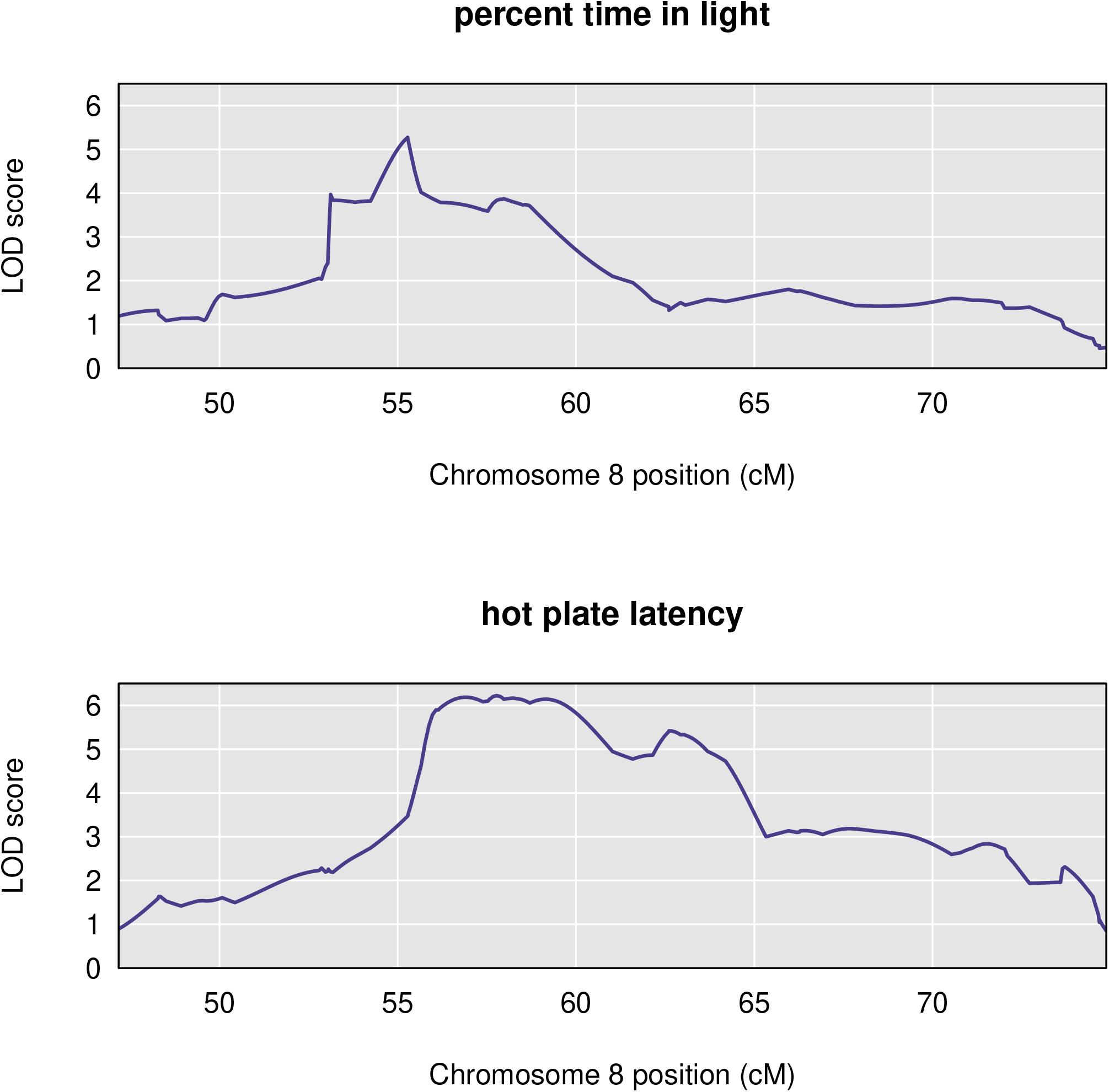
Chromosome 8 univariate LOD scores for percent time in light and hot plate latency reveal broad, overlapping peaks between 53 cM and 64 cM. The peak for percent time in light spans the region from approximately 53 cM to 60 cM, with a maximum near 55 cM. The peak for hot plate latency begins near 56 cM and ends about 64 cM.

The estimated QTL allele effects for the two traits are quite different (Figure 3). With the QTL placed at 55 cM, for “percent time in light”, the WSB and PWK alleles are associated with large phenotypes and NOD with low phenotypes. For “hot plate latency”, on the other hand, CAST and NZO show low phenotypes and NOD and PWK are near the center.

**Figure 3.**
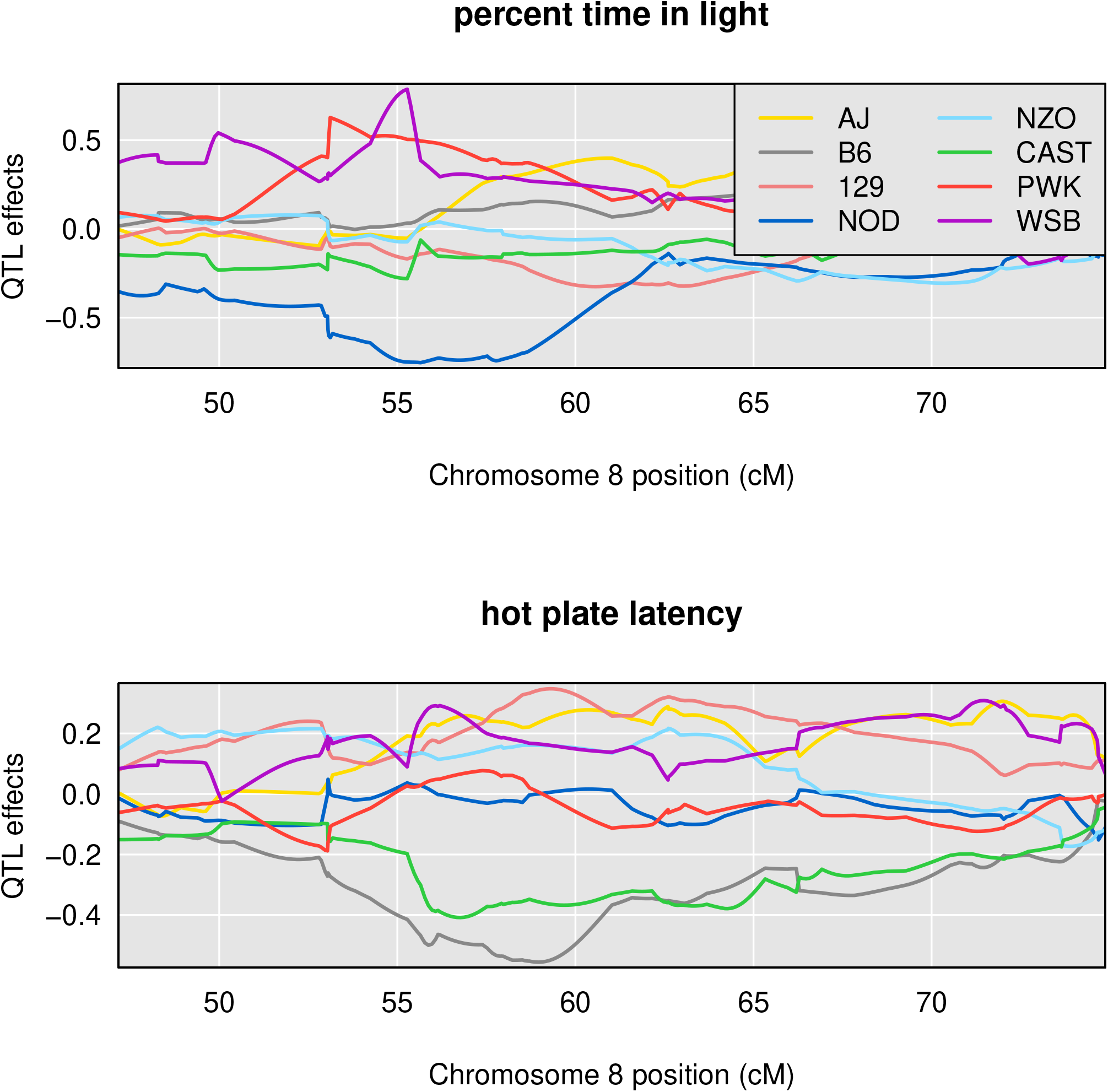
Chromosome 8 founder allele effects for percent time in light and hot plate latency demonstrate distinct allele patterns between 53cM and 64 cM.

In applying our test for pleiotropy, we performed a two-dimensional, two-QTL scan for the pair of phenotypes. With these results, we created a profile LOD plot (Figure 4). The profile LOD for “percent time in light” (in brown) peaks near 55 cM, as was seen in the univariate analysis. The profile LOD for “hot plate latency” (in blue) peaks near 57 cM, also similar to the univariate analysis. The pleiotropy trace (in gray) peaks near 55 cM.

**Figure 4.**
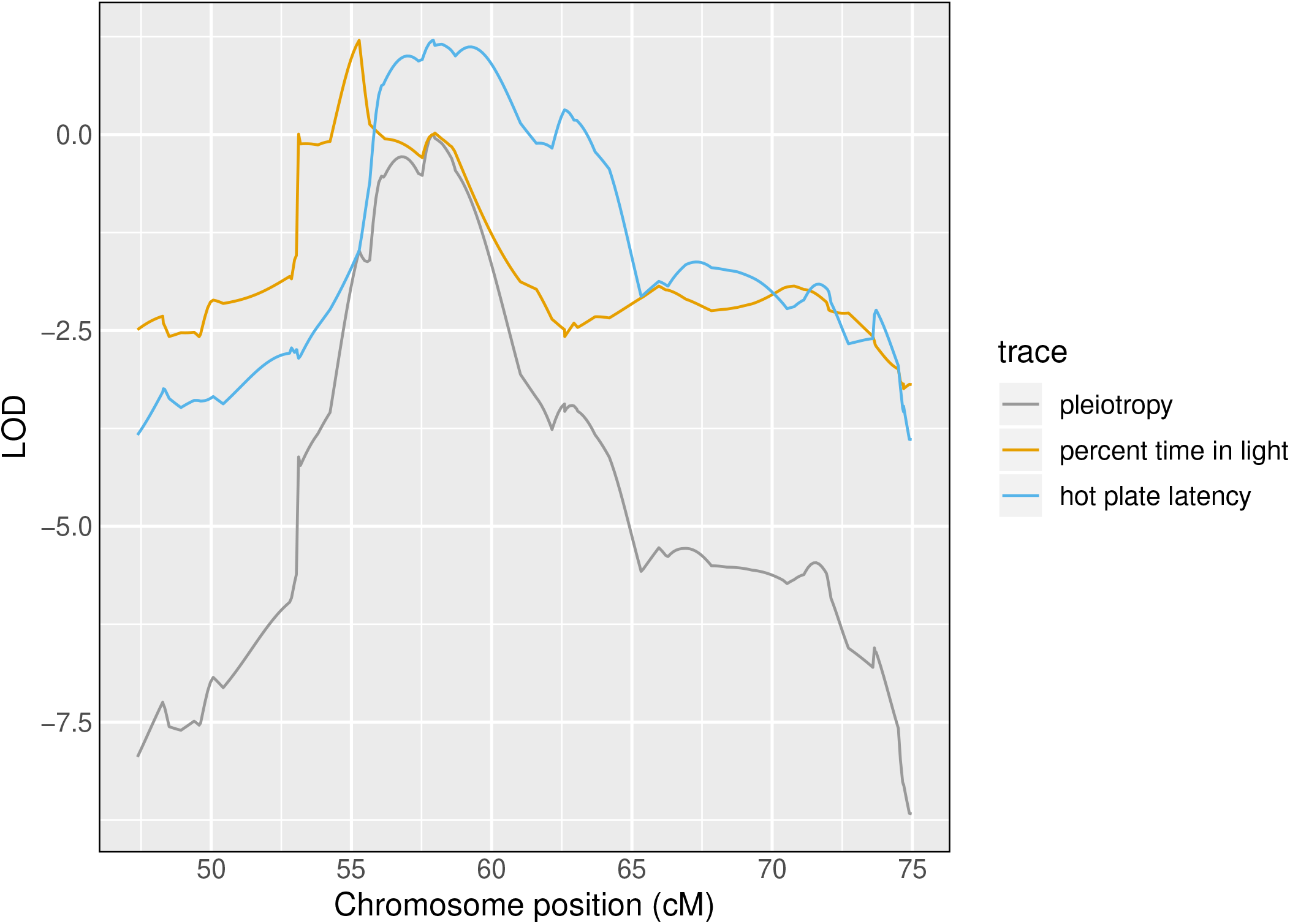
Profile LOD curves for the pleiotropy vs. separate QTL hypothesis test for “percent time in light” and “hot plate latency”. Gray trace denotes pleiotropy LOD values. Likelihood ratio test statistic value corresponds to the height of the blue and gold traces at their maxima.

The likelihood ratio test statistic for the test of pleiotropy was 1.2. Based on a parametric bootstrap with 1,000 bootstrap replicates, the estimated p-value was 0.11. Thus, by our approach, the evidence for the two traits having distinct QTL is weak.

## Discussion

We developed a test of pleiotropy vs. separate QTL for multiparental populations, extending the work of Jiang and Zeng (1995) for multiple alleles and with a linear mixed model to account for population structure (Kang *et al*. 2010; Yang *et al*. 2014). Our simulation studies indicate that our test is slightly conservative, with type I error rates below their nominal values (Table 1). Power is affected by many factors (including sample size, effect size, and allele frequencies). We studied the effects of interlocus distance and QTL effect on power, and we showed that our test has power to detect presence of separate loci, especially when univariate trait associations are strong (Figure 1). Type I error rates indicate that our test is slightly conservative (Table 1).

In the application of our method to two behavioral phenotypes in a study of 261 Diversity Outbred mice (Recla *et al*. 2014; Logan *et al*. 2013), the evidence for the presence of two distinct QTL, with one QTL (which contains the *Hydin* gene) affecting only “hot plate latency” and a second QTL affecting “percent time in light” was weak (p = 0.11, Figure 4).

Founder allele effects plots provide further evidence for the presence of two distinct loci. As Macdonald and Long (2007) and King *et al*. (2012) have demonstrated in their analyses of multi-parental *Drosophila* populations, a biallelic pleiotropic QTL would result in allele effects plots that have similar patterns. While we do not know that “percent time in light” and “hot plate latency” arise from biallelic QTL, the dramatic differences that we observe in allele effects patterns further support the argument for two distinct loci.

We have implemented our methods in an R package `qtl2pleio`, but analyses can be computationally intensive and time consuming. `qtl2pleio` is written mostly in R, and so we could likely obtain improved computational speed by porting parts of the calculations to a compiled language such as C or C++. To accelerate our multi-dimensional QTL scans, we have integrated C++ code into `qtl2pleio`, using the Rcpp package (Eddelbuettel *et al*. 2011).

Another computational bottleneck is the estimation of the variance components *V_g_* and *V_e_*. To accelerate this procedure, especially for the joint analysis of more than two traits, we will consider other strategies for variance component estimation, including that described by Meyer *et al*. (2018). Meyer *et al*. (2018), in joint analysis of dozens of traits, implement a bootstrap strategy to estimate variance components for lower-dimensional phenotypes before combining bootstrap estimates into valid covariance matrices for the full multivariate phenotype. Such an approach may ease some of the computational burdens that we encountered.

We view tests of pleiotropy as complementary to mediation tests and related methods that have become popular for inferring biomolecular causal relationships (Chick *et al*. 2016; Schadt *et al*. 2005; Baron and Kenny 1986). A mediation test proceeds by including a putative mediator as a covariate in the regression analysis of phenotype and QTL genotype; a substantial reduction in the association between genotype and phenotype corresponds to evidence of mediation.

Mediation analyses and our pleiotropy test ask distinct, but related, questions. Mediation analysis seeks to establish causal relationships among traits, including molecular traits, or dependent biological and behavioral processes. Pleiotropy tests examine whether two traits share a single source of genetic variation, which may act in parallel or in a causal network. Pleiotropy is required for causal relations among traits. In many cases, the pleiotropy hypothesis is the only reasonable one.

Schadt *et al*. (2005) argued that both pleiotropy tests and causal inference methods may contribute to gene network reconstruction. They developed a model selection strategy, based on the Akaike Information Criterion (Akaike 1974), to determine which causal model is most compatible with the observed data. Schadt *et al*. (2005) extended the methods of Jiang and Zeng (1995) to consider more complicated alternative hypotheses, such as the possibility of two QTL, one of which associates with both traits, and one of which associates with only one trait. As envisioned by Schadt *et al*. (2005), we foresee complementary roles emerging for our pleiotropy test and mediation tests in the dissection of complex trait genetic architecture.

Two related approaches for identifying and exploiting pleiotropy deserve mention. First, CAPE (Combined Analysis of Pleiotropy and Epistasis) is a strategy for identifying higher-order relationships among traits and marker genotypes (Tyler *et al*. 2013, 2016) and has recently been extended for use with multiparental populations, including DO mice (Tyler *et al*. 2017). CAPE exploits the pleiotropic relationship among traits in order to characterize the underlying network of QTLs, and it can suggest possible pleiotropic effects, but it does not provide an explicit test of pleiotropy. Second, Schaid *et al*. (2016) described a test for pleiotropy in the context of human genome-wide association studies (GWAS). Their approach is fundamentally different from ours, in that rather than ask whether traits are affected by a common locus or distinct loci, they ask whether the traits are all affected by a particular SNP or only some are. The difference in these approaches may be attributed to the difference in mapping resolution between human GWAS and experimental populations.

Technological advances in mass spectrometry and RNA sequencing have enabled the acquisition of high-dimensional biomolecular phenotypes (Ozsolak and Milos 2011; Han *et al*. 2012). Mul-tiparental populations in *Arabidopsis*, maize, wheat, oil palm, rice, *Drosophila*, yeast, and other organisms enable high-precision QTL mapping (Yu *et al*. 2008; Tisné *et al*. 2017; Stanley *et al*. 2017; Raghavan *et al*. 2017; Mackay *et al*. 2012; Kover *et al*. 2009; Cubillos *et al*. 2013). The need to analyze high-dimensional phenotypes in multiparental populations compels the scientific community to develop tools to study genotype-phenotype relationships and complex trait architecture. Our test, and its future extensions, will contribute to these ongoing efforts.

## Acknowledgments

The authors thank Lindsay Traeger, Julia Kemis, Qiongshi Lu, Rene Welch, and two anonymous referees for valuable suggestions to improve the manuscript. This work was supported in part by National Institutes of Health grants R01GM070683 (to K.W.B.) and P50DA039841 (to E.J.C.). The research made use of compute resources and assistance of the UW-Madison Center For High Throughput Computing (CHTC) in the Department of Computer Sciences at UW-Madison, which is supported by the Advanced Computing Initiative, the Wisconsin Alumni Research Foundation, the Wisconsin Institutes for Discovery, and the National Science Foundation, and is an active member of the Open Science Grid, which is supported by the National Science Foundation and the U.S. Department of Energy’s Office of Science.

---

**Table S1.** Eight founder lines and their one-letter abbreviations.

| Founder allele | One-letter abbreviation |
| --- | --- |
| A/J | A |
| C57BL/6J | B |
| 129S1/SvImJ | C |
| NOD/ShiLtJ | D |
| NZO/H1LTJ | E |
| Cast/EiJ | F |
| PWK/PhJ | G |
| WSB/EiJ | H |

**Table S2.** Both “hot plate latency” and “percent time in light” demonstrate multiple QTL peaks with LOD scores above 5.

| phenotype | chr | pos | LOD score |
| --- | --- | --- | --- |
| percent time in light | 8 | 55.28 | 5.27 |
| hot plate latency | 8 | 57.77 | 6.22 |
| percent time in light | 9 | 36.70 | 5.42 |
| hot plate latency | 9 | 46.85 | 5.22 |
| percent time in light | 11 | 63.39 | 6.46 |
| hot plate latency | 12 | 43.52 | 5.13 |
| percent time in light | 15 | 15.24 | 5.67 |
| hot plate latency | 19 | 47.80 | 5.48 |

**Figure S1.**
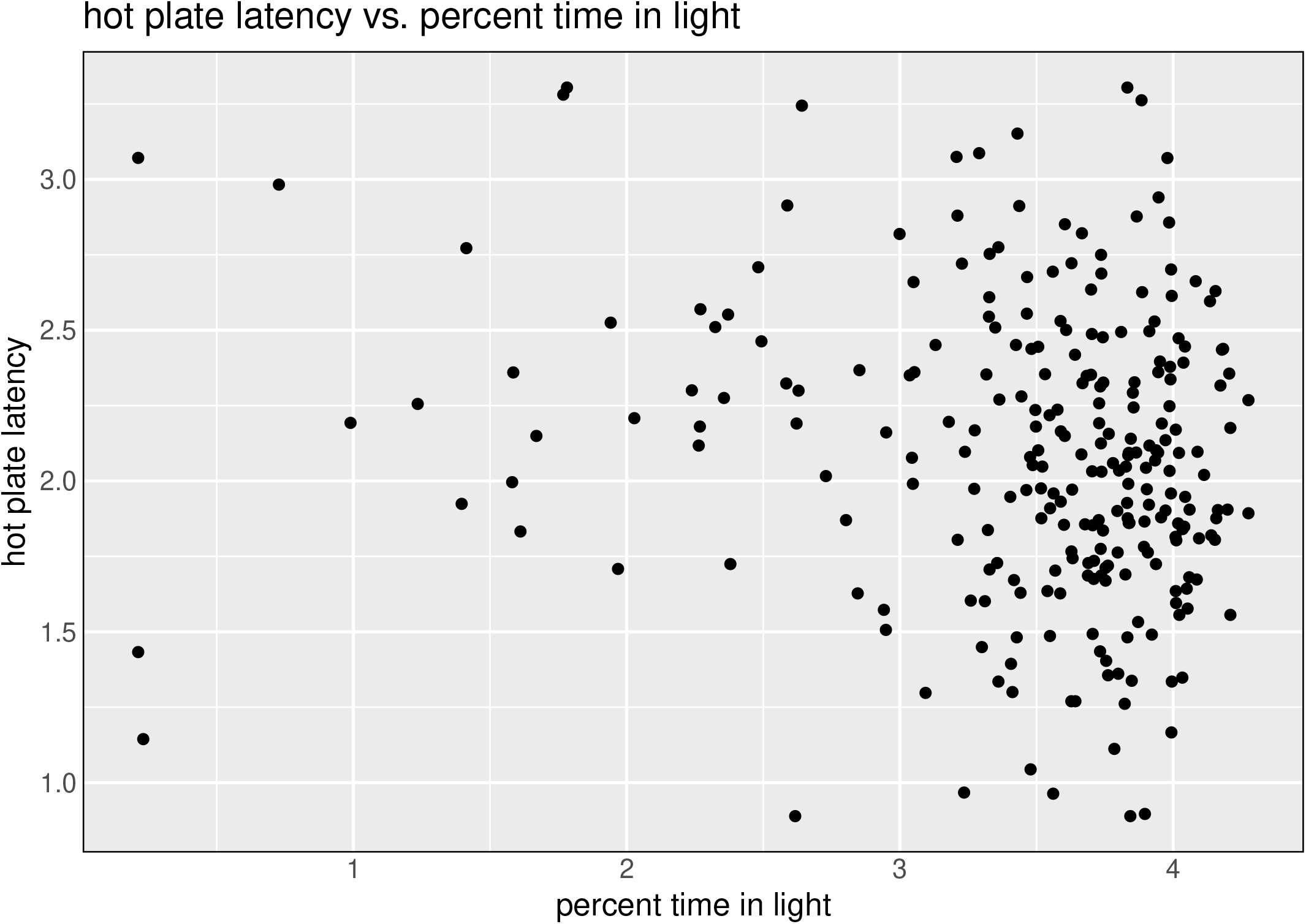
Scatter plot of “hot plate latency” against “percent time in light”, after applying logarithm transformations and winsorizing both traits.

**Figure S2.**
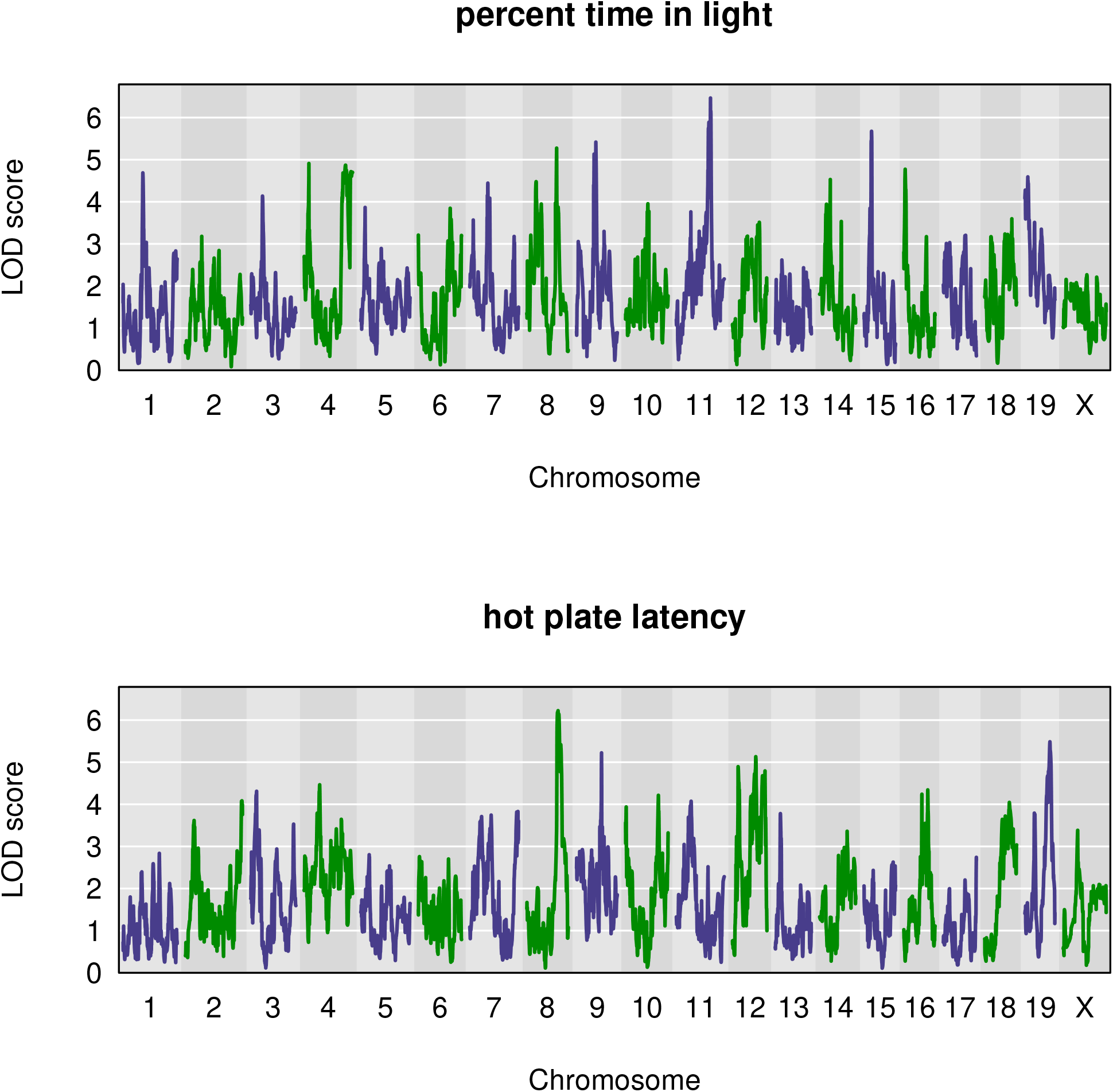
Genome-wide QTL scan for percent time in light reveals multiple QTL, including one on Chromosome 8.

**Figure S3.**
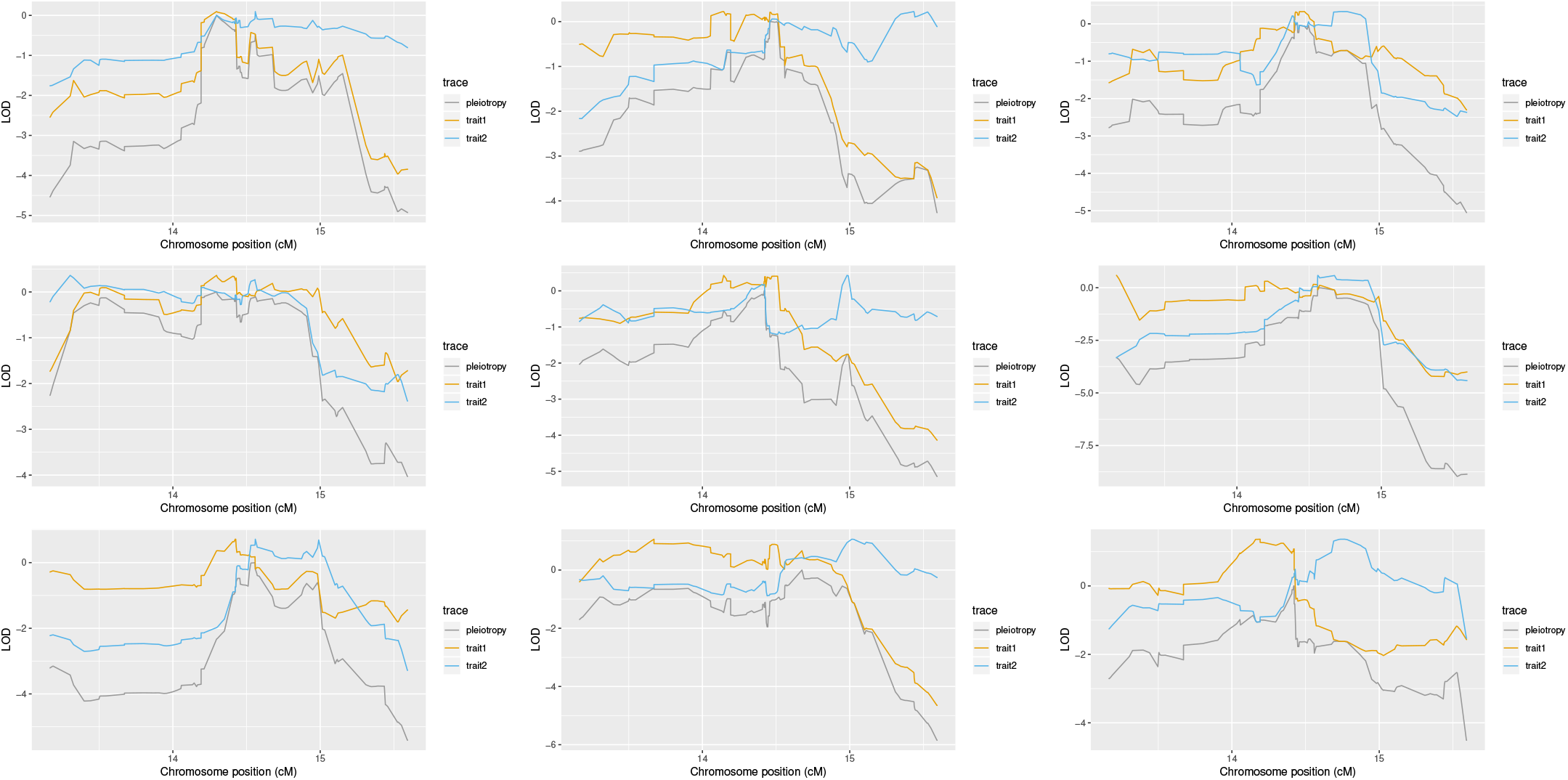
Randomly sampled profile LOD plots for trait pairs simulated with interlocus distance equal to 0.5 cM, effect size 1, and allele partitioning F:ABCDEGH. The test statistics, by row, starting at the top and proceeding left to right, are: 0.10, 0.22, 0.33, 0.36, 0.43, 0.59, 0.71, 1.06, and 1.37. The critical value (for *α* = 0.05) is 0.77.

## Literature Cited

Akaike, H., 1974 A new look at the statistical model identification. IEEE T. Automat. Contr. 19: 716–723.

Baron, R. M. and D. A. Kenny, 1986 The moderator–mediator variable distinction in social psychological research: Conceptual, strategic, and statistical considerations. J. Pers. Soc. Psychol. 51: 1173–1182.

Boehm, F., 2018 gemma2: GEMMA multivariate linear mixed model. R package version 0.0.1.

Broman, K. W., D. M. Gatti, P. Simecek, N. A. Furlotte, P. Prins, et al., 2019 R/qtl2: Software for mapping quantitative trait loci with high-dimensional data and multi-parent populations. Genetics 211: 495–502.

Chesler, E. J., D. M. Gatti, A. P. Morgan, M. Strobel, L. Trepanier, et al., 2016 Diversity outbred mice at 21: Maintaining allelic variation in the face of selection. G3 6: 3893–3902.

Chick, J. M., S. C. Munger, P. Simecek, E. L. Huttlin, K. Choi, et al., 2016 Defining the consequences of genetic variation on a proteome-wide scale. Nature 534: 500–505.

Churchill, G. A., D. C. Airey, H. Allayee, J. M. Angel, A. D. Attie, et al., 2004 The Collaborative Cross, a community resource for the genetic analysis of complex traits. Nat. Genet. 36: 1133–1137.

Churchill, G. A., D. M. Gatti, S. C. Munger, and K. L. Svenson, 2012 The diversity outbred mouse population. Mamm. Genome 23: 713–718.

Cubillos, F. A., L. Parts, F. Salinas, A. Bergström, E. Scovacricchi, et al., 2013 High-resolution mapping of complex traits with a four-parent advanced intercross yeast population. Genetics 195: 1141–1155.

de Koning, D.-J. and L. M. McIntyre, 2014 Genetics and G3: Community-driven science, community-driven journals. Genetics 198: 1–2.

Eddelbuettel, D., R. François, J. Allaire, J. Chambers, D. Bates, et al., 2011 Rcpp: Seamless R and C++ integration. J. Stat. Softw. 40: 1–18.

Efron, B., 1979 Bootstrap methods: another look at the jackknife. Ann. Stat. 7: 1–26.

Han, X., K. Yang, and R. W. Gross, 2012 Multi-dimensional mass spectrometry-based shotgun lipidomics and novel strategies for lipidomic analyses. Mass Spectrom. Rev. 31: 134–178.

Jiang, C. and Z.-B. Zeng, 1995 Multiple trait analysis of genetic mapping for quantitative trait loci. Genetics 140: 1111–1127.

Kang, H. M., J. H. Sul, S. K. Service, N. A. Zaitlen, S.-y. Kong, et al., 2010 Variance component model to account for sample structure in genome-wide association studies. Nat. Genet. 42: 348–354.

Keller, M. P., D. M. Gatti, K. L. Schueler, M. E. Rabaglia, D. S. Stapleton, et al., 2018 Genetic drivers of pancreatic islet function. Genetics 209: 335–356.

King, E. G., C. M. Merkes, C. L. McNeil, S. R. Hoofer, S. Sen, et al., 2012 Genetic dissection of a model complex trait using the Drosophila Synthetic Population Resource. Genome Res. 22: 1558–1566.

Knott, S. A. and C. S. Haley, 2000 Multitrait least squares for quantitative trait loci detection. Genetics 156: 899–911.

Kover, P. X., W. Valdar, J. Trakalo, N. Scarcelli, I. M. Ehrenreich, et al., 2009 A multiparent advanced generation inter-cross to fine-map quantitative traits in *Arabidopsis thaliana*. PLoS Genet. 5: e1000551.

Logan, R. W., R. F. Robledo, J. M. Recla, V. M. Philip, J. A. Bubier, et al., 2013 High-precision genetic mapping of behavioral traits in the diversity outbred mouse population. Genes Brain Behav. 12: 424–437.

Macdonald, S. J. and A. D. Long, 2007 Joint estimates of QTL effect and frequency using synthetic recombinant populations of *Drosophila melanogaster*. Genetics 176: 1261–1281.

Mackay, T. F., S. Richards, E. A. Stone, A. Barbadilla, J. F. Ayroles, et al., 2012 The *Drosophila melanogaster* genetic reference panel. Nature 482: 173–178.

Meyer, H. V., F. P. Casale, O. Stegle, and E. Birney, 2018 LiMMBo: a simple, scalable approach for linear mixed models in high-dimensional genetic association studies. bioRxiv. DOI: https://doi.org/10.1101/255497.

Meyer, K., 1989 Restricted maximum likelihood to estimate variance components for animal models with several random effects using a derivative-free algorithm. Genet. Sel. Evol. 21: 317–340.

Meyer, K., 1991 Estimating variances and covariances for multivariate animal models by restricted maximum likelihood. Genet. Sel. Evol. 23: 67–83.

Ozsolak, F. and P. M. Milos, 2011 RNA sequencing: advances, challenges and opportunities. Nat. Rev. Genet. 12: 87–98.

Raghavan, C., R. Mauleon, V. Lacorte, M. Jubay, H. Zaw, et al., 2017 Approaches in characterizing genetic structure and mapping in a rice multiparental population. G3 7: 1721–1730.

Recla, J. M., R. F. Robledo, D. M. Gatti, C. J. Bult, G. A. Churchill, et al., 2014 Precise genetic mapping and integrative bioinformatics in Diversity Outbred mice reveals *Hydin* as a novel pain gene. Mamm. Genome 25: 211–222.

Schadt, E. E., J. Lamb, X. Yang, J. Zhu, S. Edwards, et al., 2005 An integrative genomics approach to infer causal associations between gene expression and disease. Nat. Genet. 37: 710–717.

Schaid, D. J., X. Tong, B. Larrabee, R. B. Kennedy, G. A. Poland, et al., 2016 Statistical methods for testing genetic pleiotropy. Genetics 204: 483–497.

Stanley, P. D., E. Ng’oma, S. O’Day, and E. G. King, 2017 Genetic dissection of nutrition-induced plasticity in insulin/insulin-like growth factor signaling and median life span in a Drosophila multiparent population. Genetics 206: 587–602.

Tian, J., M. P. Keller, A. T. Broman, C. Kendziorski, B. S. Yandell, et al., 2016 The dissection of expression quantitative trait locus hotspots. Genetics 202: 1563–1574.

Tisné, S., V. Pomiès, V. Riou, I. Syahputra, B. Cochard, et al., 2017 Identification of ganoderma disease resistance loci using natural field infection of an oil palm multiparental population. G3 7: 1683–1692.

Tyler, A. L., L. R. Donahue, G. A. Churchill, and G. W. Carter, 2016 Weak epistasis generally stabilizes phenotypes in a mouse intercross. PLoS Genet. 12: e1005805.

Tyler, A. L., B. Ji, D. M. Gatti, S. C. Munger, G. A. Churchill, et al., 2017 Epistatic networks jointly influence phenotypes related to metabolic disease and gene expression in diversity outbred mice. Genetics 206: 621–639.

Tyler, A. L., W. Lu, J. J. Hendrick, V. M. Philip, and G. W. Carter, 2013 CAPE: an R package for combined analysis of pleiotropy and epistasis. PLoS Comput. Biol. 9: e1003270.

Yang, J., N. A. Zaitlen, M. E. Goddard, P. M. Visscher, and A. L. Price, 2014 Advantages and pitfalls in the application of mixed-model association methods. Nat. Genet. 46: 100–106.

Yu, J., J. B. Holland, M. D. McMullen, and E. S. Buckler, 2008 Genetic design and statistical power of nested association mapping in maize. Genetics 178: 539–551.

Zeng, Z.-B., J. Liu, L. F. Stam, C.-H. Kao, J. M. Mercer, et al., 2000 Genetic architecture of a morphological shape difference between two Drosophila species. Genetics 154: 299–310.

Zhou, X. and M. Stephens, 2014 Efficient multivariate linear mixed model algorithms for genome-wide association studies. Nat. Methods 11: 407–409.

